# Species composition of Culicidae (Diptera) in Bromeliads in rural and urban areas of Londrina, Brazil

**DOI:** 10.1101/2023.05.20.538492

**Authors:** Bianca Piraccini Silva, Adriano Nobre Arcos, Francisco Augusto da Silva Ferreira, Cristiano Medri, João Antonio Cyrino Zequi

## Abstract

Bromeliaceae, has 82 genera and 3,719 species found throughout the Neotropical region, from the southern United States to central Argentina and Chile. These plants are present in environments with different characteristics and have high endemism in the Atlantic Forest in Brazil. Tank bromeliads stand out for their morphology, which forms permanent water-accumulating compartments called phytotelmata, rich in organic detritus and living organisms. Culicidae use phytotelma as a breeding ground, and some species of Wyeomyia are highly dependent on these tanks. Research on the importance of bromeliads as water reservoirs that may favor the reproduction of mosquitoes that carry pathogens are controversial. Some studies indicate that these plants are potential breeding grounds for *Aedes*, while others suggest that they are not preferential foci for synanthropic mosquitoes, as is the case for *A. aegypti*. This work aimed at verifying the presence of *Aedes* and other Culicidae in native bromeliads in natural habitat and under ornamental cultivation in an urban environment, comparing the incidence of larvae in these possible breeding sites and contrasting with the luminosity and accumulated water volumes. Differences in species composition were observed between rural and urban environments. In the rural environment, four species of Culicidae were collected, the most abundant being *Wyeomyia galvaoi* -19 specimens, followed by *Culex* (*Microculex*) *imitator* - five specimens, while *Aedes aegypti* and *Toxorhynchites* sp 1, were the least abundant, with two individuals each. In the urban environment, five species were found, with greater abundance of *Aedes aegypti* (461 individuals), followed by *Aedes albopictus* (45), *Toxorhynchites* sp. 01 (3), *Wyeomyia* sp1 (02), and *Wyeomyia galvaoi* (01) In addition, these phytotelma showed great diversity of other organisms, such as: mites, nematodes, protists, rotifers, ostracods, and insect larvae of the Chironomidae, Syrphidae and Tachinidae families. The study concluded that the incidence of mosquito larvae in native bromeliads is low, and that luminosity and accumulated water volumes can influence the presence of larvae. The authors emphasize that the importance of bromeliads in mosquito reproduction may vary according to several factors.

## INTRODUCTION

The Bromeliaceae family comprises 82 genera and 3719 species, widely dispersed in the Neotropical region, extending from the southern United States to central Argentina and Chile. These plants are present in environments with different characteristics, from extremely dry and sunny to humid and shady, and at different altitudes and temperatures (Benzing, 2000; Gouda & Butcher, 2022). Only one species, *Pitcairnia feliciana* (A.Chev.) Harms & Mildbr., is found on the African continent.

These plants are present in all Brazilian biomes, especially in the Atlantic Forest, where they represent one of the most relevant taxonomic groups, due to the high degree of endemism. The Bromeliaceae family has important ecological value, mainly due to its interactions with fauna, constituting a micro-habitat for several species of vertebrates and invertebrates (Martinelli et al. 2008; Docile et al. 2017).

In this context, the tank bromeliads stand out, presenting ecological interactions that occur due to the morphology of the species, which are endowed with rosette phyllotaxis and arranged leaf sheaths, forming permanent water accumulator compartments called phytotelmata, that are rich in organic debris and living organisms (Benzing, 2000). The water accumulated in the plants is used as a water trough, foraging site, for shelter, and refuge from predators, as well as for breeding and nesting sites (Paula, 2004). It is also noteworthy that these tanks are places of oviposition and larval development for several orders of insects, including mosquitoes of the Culicidae family.

Culicidae use phytotelma as a breeding ground, in this way the accumulated water is utilized as a place for the development of their immature forms. Some genera even have a high dependence on these tanks, such as some species of *Wyeomyia* (Frank & Lounibos, 2009). The importance of bromeliads as water reservoirs that can favor the reproduction of mosquito vectors of pathogens is controversial. Research has produced contradictory results on the findings of immature forms of *Aedes aegypti* Linnaeus and *Aedes albopictus* Skuse in their phytotelma.

Some studies point out that these plants are potential breeding grounds for *Aedes* (Natal et al. 1997; Forattini et al. 1998; Forattini & Marques 2000; Marques et al. 2001; Cunha et al. 2002; Ceretti-Junior et al. 2016; Wilke et al. 2018). On the other hand, some findings indicate that bromeliads are not the preferential foci of synanthropic mosquitoes, as is the case with *A. aegypti* (Ospina-Batista et al. 2008; Frank & Lounibos 2009; Lopez et al. 2009; Mocellin 2010; Santos et al. 2011; Oliveira & Almeida-Neto 2017). The authors emphasize that bromeliads form a complex scenario, in which their importance in mosquito reproduction can vary according to the location, habitat, climate, biotic and abiotic characteristics of the phytotelma, and species of vector and bromeliad, in addition to human behavior.

Given the above, the current work sought to verify the presence of *Aedes* and other Culicidae in native bromeliads occurring in the natural habitat and under ornamental cultivation in an urban environment; comparing the incidence of larvae in these possible breeding sites and contrasting with the luminosity and accumulated volumes of water; seeking to answer if, and under what conditions, these plants can be breeding grounds for *Aedes* larvae and other Culicidae.

## MATERIAL AND METHODS

### Study area

The study was conducted in two locations in the city of Londrina, Paraná state, southern Brazil, a region with a Cfa (subtropical) climate, according to the Köppen classification, with an average temperature in the coldest month of below 18°C and average in the hottest month of above 22°C (Wrege et al. 2011).

The first location (23° 38’ 17.44” S 51° 5’ 12.48” W; with 711 meters) (Google Earth LCC) is in a rural area with basaltic outcrops covered by rupicolous and endemic vegetation, composed, in addition to other species and families of the bromeliads *Dyckia walteriana* Leme, *Dyckia leptostachya* Baker, and *Bromelia balansae* Mez., as well as the tank bromeliad *Aechmea distichantha* Lem. The second location is in a residence in an urban area (23° 18’ 49.13”S, 51° 11’ 24.51 ””W; with 553 meters) (Google Earth LCC); containing a sunny part of the garden consisting of a green roof and a ground floor area, with cultivation sites for the tank bromeliads *Alcantarea imperialis* (Carriere) Harms, *Aechmea blanchetiana* (Baker) LB Sm. var. rubra, *A. blanchetiana* var. limão, *A. distichantha, Aechmea pineliana* (Brong. Planch.) Baker., and *Neoregelia cruenta* (R.Graham) LB Sm.; and a shady part of the garden composed of a sloping area and a flat area, places where the following tank bromeliads are cultivated *A. blanchetiana* var. rubra, *Aechmea nudicaulis* (L.), *Neoregelia sp*. (híbrida), and *Vriesea guttata Linden & André*. Of all the species cultivated in the residence in the urban area, only *A. distichantha* is native to the region of Londrina, PR (Figure 1).

**Figure 1.**
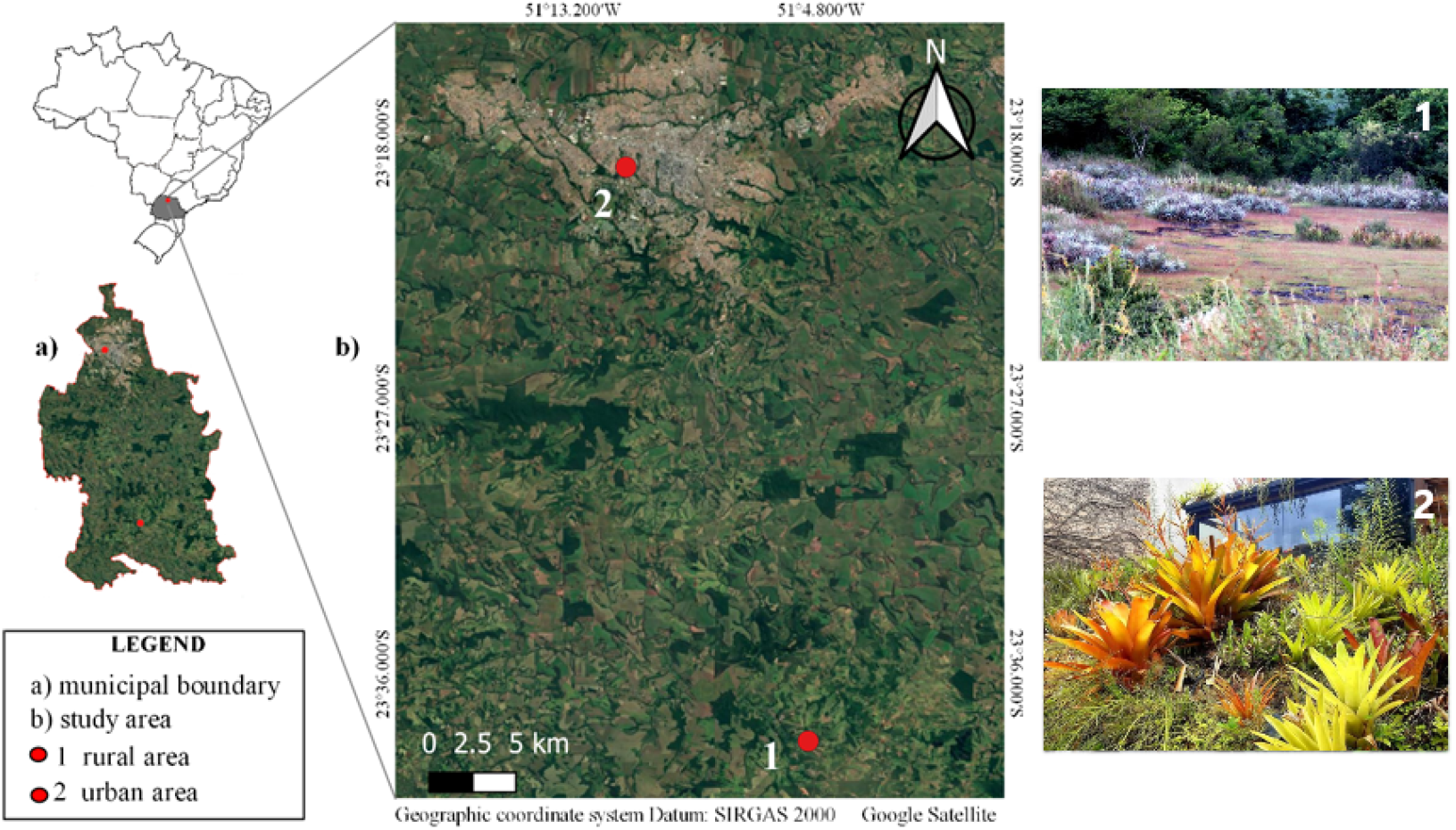
Geographic location and images of the locations of *Aedes* collection points in the rural and urban areas of the municipality of Londrina, state of Paraná, Brazil, between March and April 2017.

**Figure 1.**
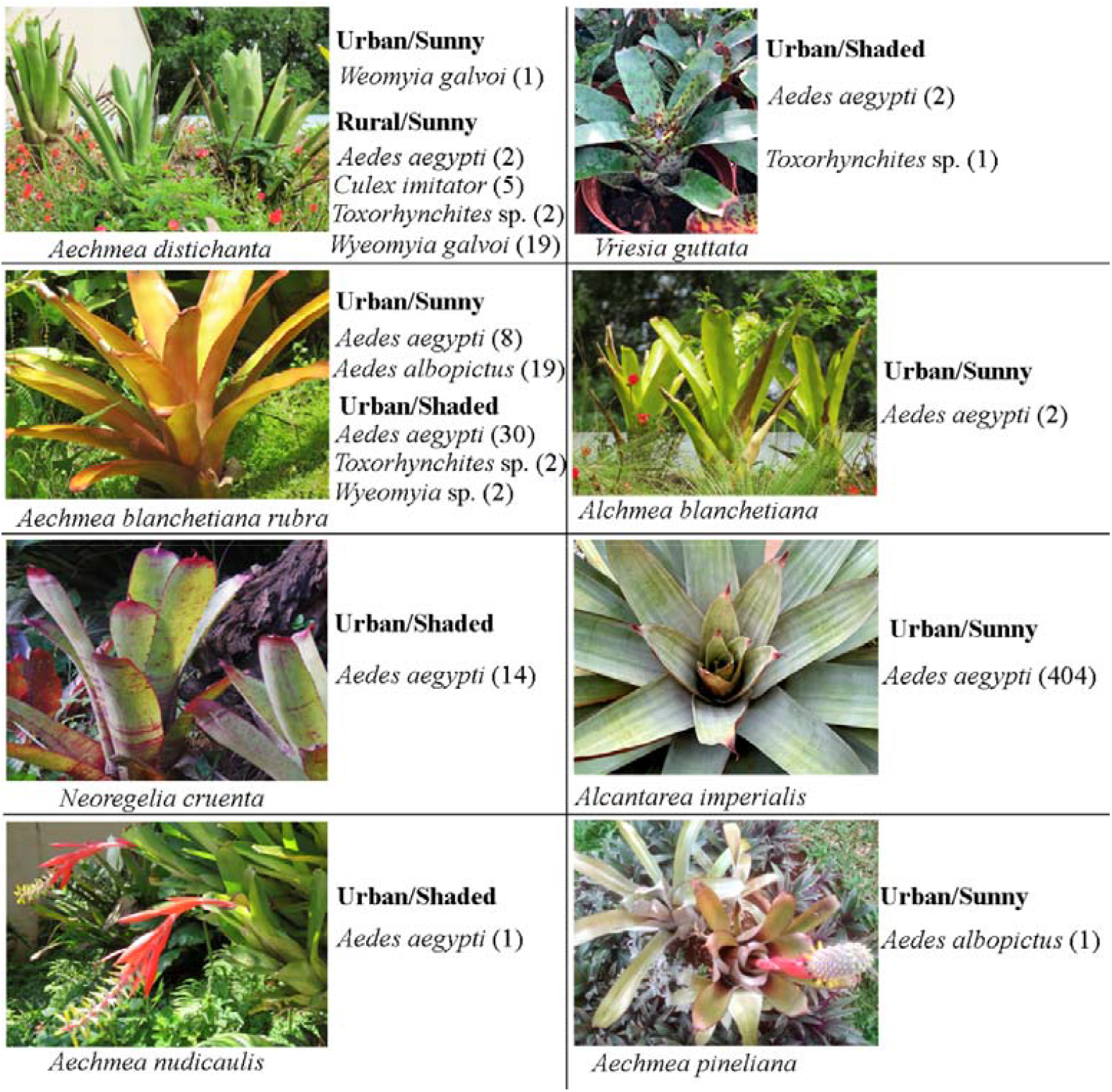
Abundance and diversity of Culicidae collected in an urban and rural environment in the municipality of Londrina, PR, between March and April 2017 in different bromeliads exposed to Sunny and Shaded areas.

### Material collection

Sampling was carried out in two locations between March and April 2017, weekly, for five consecutive weeks. Thermo-hygrometers were installed at both locations to measure the temperature (°C) and relative humidity (%). In addition, the rainfall data measured by the Londrina meteorological station during the experimental period were obtained from the Paraná Meteorological System (SIMEPAR).

As a way of monitoring the presence of *Aedes*, traps to capture eggs (ovitraps) were installed in each location. At point 1, five individuals of *Aechmea distichantha* were monitored, and at point 2, five individuals of each species/environment (sun or shade) mentioned above, with the exception of *Alcantarea imperialis*, for which three individuals were sampled as this was the total number available.

During this period, water from the bromeliads was extracted with the aid of a silicone tube attached to a syringe. The life forms found were collected with a sieve (1mm mesh) and the volume of water was measured using a beaker. The water was later returned to the tank bromeliads. The collected life forms were taken for identification and counting with the aid of optical microscopy, together with samples of the water accumulated in each bromeliad, to the Laboratory of General and Medical Entomology of the State University of Londrina. Identification of the taxonomic groups collected was carried out, as well as the number of individuals collected in each group, with emphasis on the larval forms of *A. aegypti* and *A. albopictus*, in addition to possible other Culicidae. The water accumulated in the bromeliads was analyzed by measuring pH, conductivity, and temperature.

### Data analysis

The data obtained for *Aechmea distichantha* were compared between the two collection points. At point 2, data were compared between sunny and shaded environments, between species, and between plants that accumulate more than one liter of water with those that accumulate less than one liter. For both collection points, the data obtained were tested in relation to the correlation between the physical-chemical water and meteorological variables during the experimental period.

Multiple linear regression was used to verify the relationship between the dependent variable (abundance of mosquitoes) and the independent variables (volume, pH, conductivity). The relationships between the larval composition of mosquitoes and environmental variables were evaluated by means of exploratory canonical redundancy analysis (CRA) (Legendre & Legendre, 1998) using the Monte Carlo test, with 9999 unrestricted permutations to determine the significance level of the variables (Lourenço, 2016).

The CRA, with only the selected variables, served as a model for the analysis of variance partition (Bocard et al. 1992), simulated with a Venn diagram. This analysis indicates how much of the variation in species composition is explained by the different components and by the interaction between the components (Peres-Neto et al. 2006) within the model. All analyses are available in the *Vegan* package and performed in R (Oksanen et al. 2011; R Development Core Team 2016).

## RESULTS

Differences in species composition were observed between rural and urban environments. In the rural environment, four species of Culicidae were collected, the most abundant being *Wyeomyia galvaoi* (Corrêa & Ramalho, 1956) with 19 specimens, followed by *Culex* (*Microculex*) *imitator* Theobald, 1903 with five specimens, while *Aedes aegypti* (Linnaeus, 1762) and *Toxorhynchites* sp 1, were the least abundant, with two individuals each. On the other hand, in the urban environment, five species were found, with greater abundance of *Aedes aegypti* (461 individuals), followed by *Aedes albopictus* (Skuse, 1894) (45), *Toxorhynchites* sp. 01 (3), *Wyeomyia* sp1 (02), and *Wyeomyia galvaoi* (01) (Figure 1). In addition, these phytotelma showed great diversity of other organisms, such as: mites, nematodes, protists, rotifers, ostracods, and insect larvae of the Chironomidae, Syrphidae, and Tachinidae families.

Multiple linear regression analysis revealed that the abundance of Culicidae in bromeliads was significantly correlated with the volume of water in the phytotelma (t-value= 5.083; p-value= 0.00143) (Figure 2). However, limnological characteristics such as conductivity and pH were not correlated with the abundance of mosquitoes (t-value= -1.288; p-value= 0.23865; t-value= -0.385; p-value= 0.71139, respectively), and had no influence on the abundance of mosquitoes.

**Figure 2.**
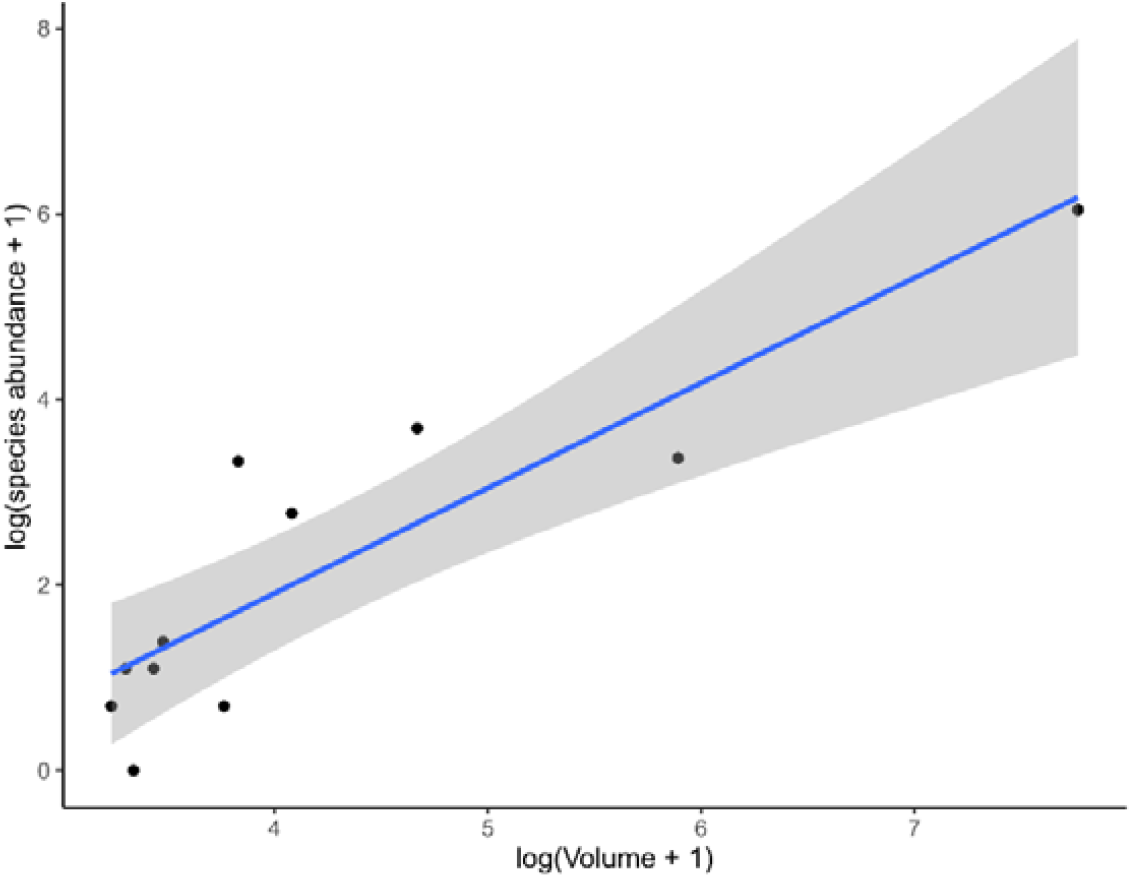
Multiple linear regression analysis of the relationship between the response variable (abundance of mosquitoes) and the explanatory variable (water volume) in bromeliads collected in urban and rural environments in the city of Londrina, PR between March and April 2017.

The environmental variables tested explained 97.3% of all the variation in the Culicidae community in bromeliads, indicating that the patterns of the larval composition of mosquitoes are greatly influenced by the environmental variables studied. The variance partition analysis showed that the water volume in the bromeliads provided an isolated explanation of 46.3 % for the mosquito species composition (Figure 3).

**Figure 3.**
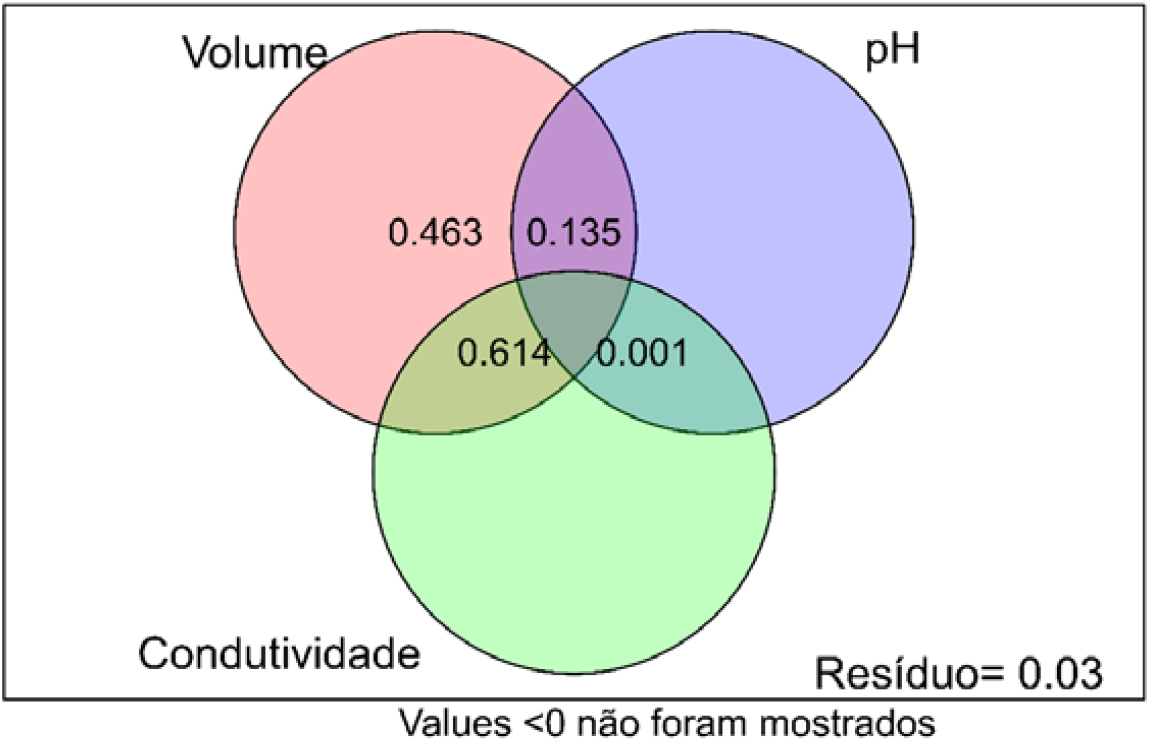
Venn diagram showing the fractions of the variance partition analysis using the matrix of environmental variables.

A positive relationship between the abundance of *A. aegypti* and bromeliads that have large volumes of water was also observed, as shown in the diagram in Figure 4.

**Figure 4.**
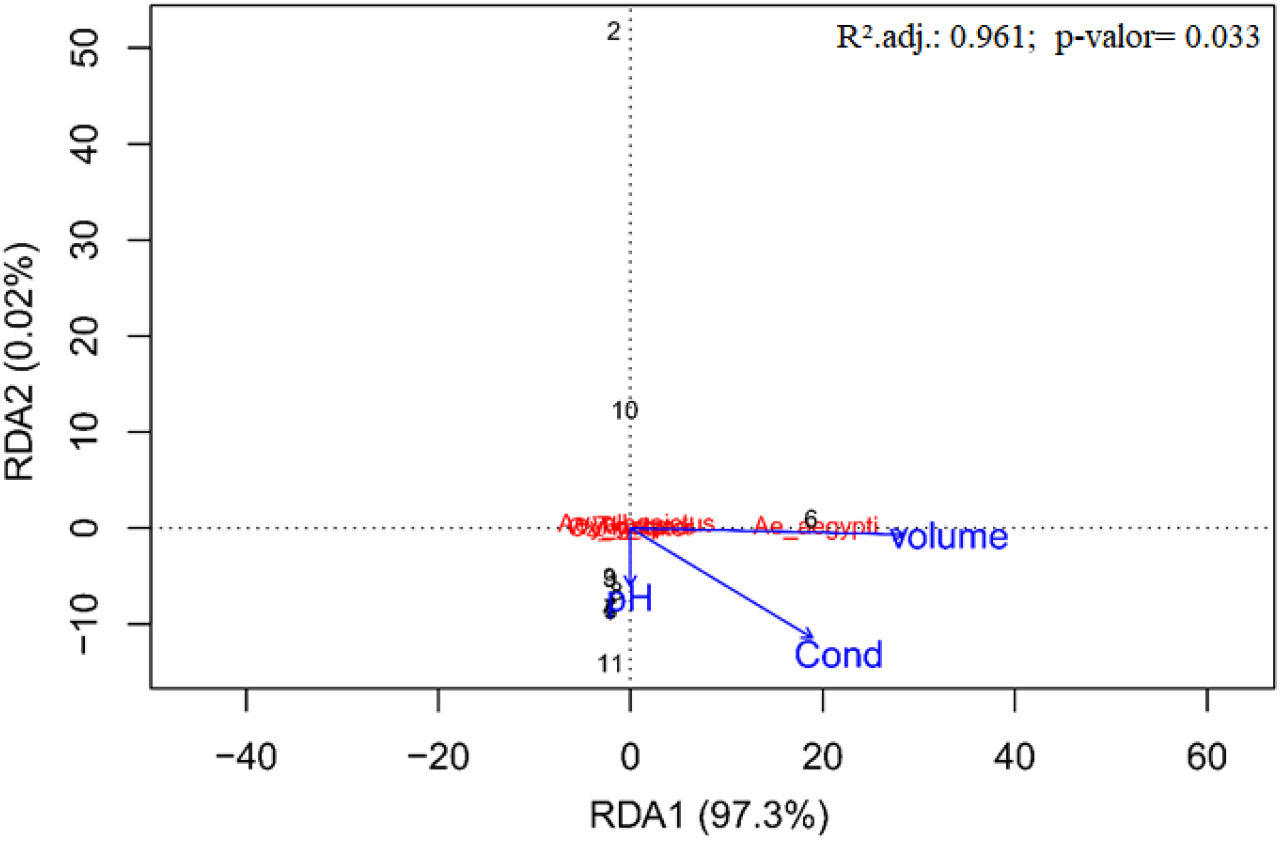
Order diagram of the redundancy analysis between the composition of mosquito species and environmental variables: volume, conductivity, and pH, in bromeliads studied in the municipality of Londrina, Paraná.

The indices of ovitrap positivity and egg density are shown in table 01. The presence of *A. aegypti* was observed in both environments, but with a much higher prevalence in the urbanized environment.

**Table 01.**
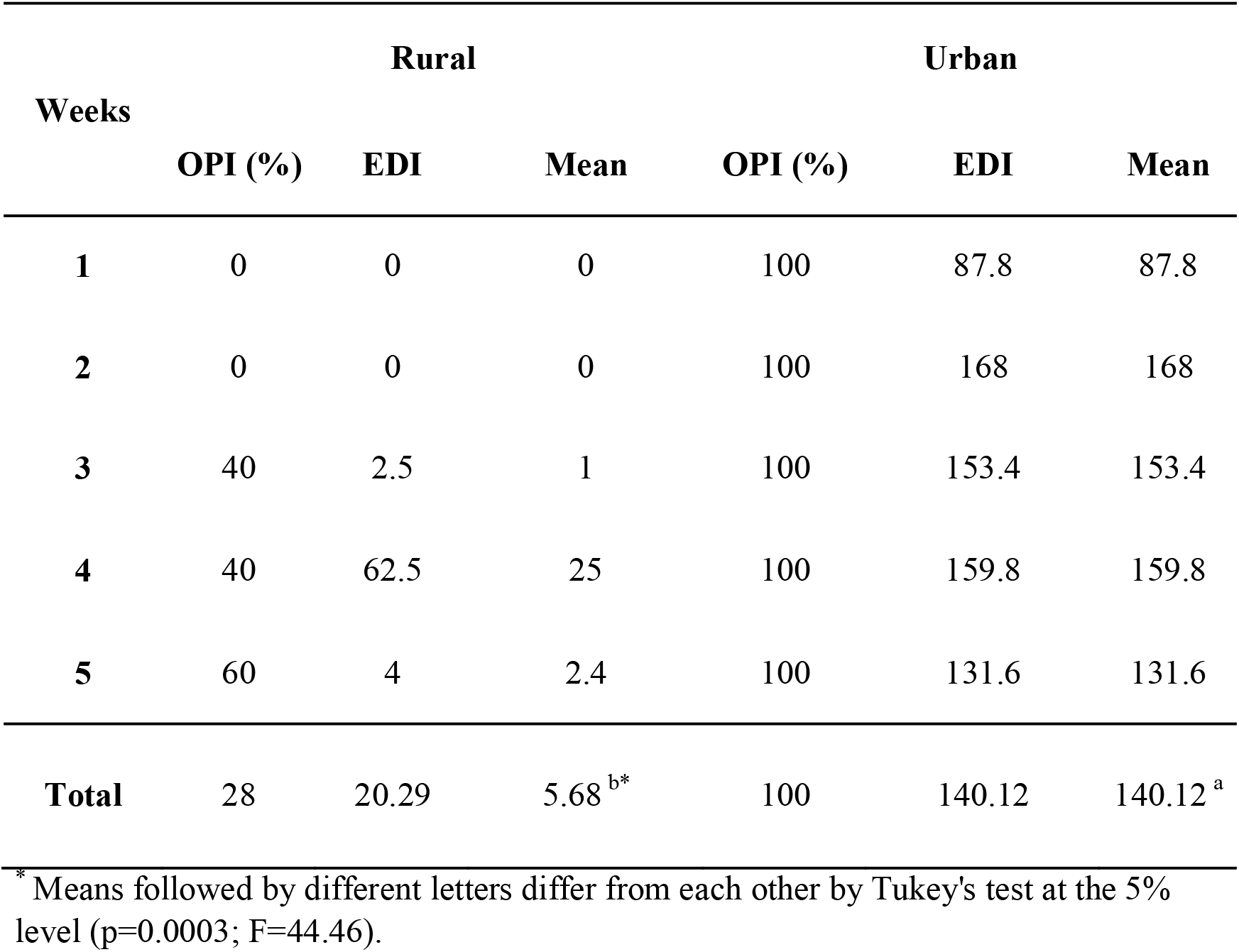
Ovitrap positivity index (OPI) and egg density index (EDI) obtained during five consecutive weeks in rural and urban areas in the city of Londrina, PR.

## DISCUSSION

Brazil experiences dengue outbreaks seasonally every year, mainly in spring and summer in colder regions. Currently, 757,068 cases have been recorded up to the eighteenth epidemiological week, covering the interval between January and May 2022. The South and Southeast regions of the country stand out as the areas with the highest occurrence of cases, with rates of 1,171 cases/100,000 inhabitants and 635.6 cases/100 thousand inhabitants, respectively.

Some works indicate that these plants are potential breeding grounds for *Aedes* (Natal et al. 1997; Forattini et al. 1998; Forattini & Marques 2000; Marques et al. 2001; Cunha et al. 2002). Wilke et al. (2018), studying ornamental tank bromeliads grown in homes and public gardens at 51 sites in Miami County, USA, suggested that ornamental bromeliads are contributing to the proliferation of *A. aegypti*, highlighting that tank bromeliads should be considered in future vector control strategies for dengue, zika, and other arboviruses. A recent study in 65 urban municipal parks in the city of São Paulo, Brazil, detected that immature forms of Culicidae, such as *A. albopictus* and *A. aegypti*, are common in bromeliad reservoirs that vegetate in these environments (Ceretti-Junior et al. 2016).

On the other hand, some studies indicate that bromeliads are not the preferred foci of *Aedes* (Mocellin 2010). According to Ospina-Batista et al. (2008), the environment of tank bromeliads makes it difficult to establish potential invaders from surrounding freshwater habitats. The differences in pH between bromeliad species are due to the decomposition processes of the accumulated organic material through release of organic acids and carbon dioxide. Lopez et al. (2009, 2011) concluded that bromeliads are the least suitable environments for the development of *A. aegypti* when compared to artificial containers, due to their acidic conditions. In addition, according to Frank & Lounibos (2009), bromeliads form a complex scenario, in which their importance in the reproduction of arbovirus vectors and other pathogens can vary according to the location, habitats, vector species, human behavior, and climate.

According to Oliveira & Almeida-Neto (2017), when studying bromeliads native to Brazil cultivated in the Jardim Botânico de Bauru, SP, the phytotelma did not constitute the foci of *A. aegypti* or *A. albopictus*, with the prevailing presence of *Culex*. The authors explained this result by the location of the Botanical Garden, far from residential areas. However, they warned that tank bromeliads grown in urban areas, owing to the reduction in artificial breeding sites, can serve as alternative breeding sites for *A. aegypti* or *A. albopictus*, and that there is a need for monitoring and entomological control in these plants.

It is generally observed that studies carried out in urbanized environments tend to point to bromeliads as possible breeding grounds for *A. aegypti*, while reports of larvae collection in phytotelma in natural environments do not find this mosquito in abundance.

The data obtained by ovitraps in the two environments studied here indicate the presence of *A. aegypti* in both environments, however in the rural environment this mosquito did not use bromeliads as a breeding ground, unlike the urban environment where a great abundance of *Aedes* was observed, especially if the plants have a large volume of water.

It is worth mentioning that the ovitraps are the most sensitive method to detect the presence of *Aedes*. In this way, it appears that bromeliads, in their natural environment, cannot be considered a breeding ground for the larvae of this mosquito, but in an urbanized environment these plants, with storage of a large volume of water, can present an important source for mosquito larvae. The importance of weekly monitoring of this phytotelma in synanthropic environments is highlighted, and if necessary, the use of mainly larvicidal products to control larvae selectively, such as *Bacillus thuringiensis* subesp. *israelenses*.

